# *Plasmodium falciparum* GBP2 is a telomere-associated protein that binds to G-quadruplex DNA and RNA

**DOI:** 10.1101/2021.07.02.450898

**Authors:** James Edwards-Smallbone, Anders L. Jensen, Lydia E. Roberts, Francis Isidore G. Totanes, Sarah R. Hart, Catherine J. Merrick

## Abstract

In the early-diverging protozoan parasite *Plasmodium*, few telomere-binding proteins have been identified and several are unique. *Plasmodium* telomeres, like those of most eukaryotes, contain guanine-rich repeats that can form G-quadruplex structures. In model systems, quadruplex-binding drugs can disrupt telomere maintenance and some quadruplex-binding drugs are potent anti-plasmodial agents. Therefore, telomere-interacting and quadruplex-interacting proteins may offer new targets for anti-malarial therapy. Here, we report that *P. falciparum* GBP2 is such a protein. It was identified via ‘Proteomics of Isolated Chromatin fragments’, applied here for the first time in *Plasmodium. In vitro, Pf*GBP2 binds specifically to G-rich telomere repeats in quadruplex form and it can also bind to G-rich RNA. *In vivo, Pf*GBP2 partially colocalises with the known telomeric protein HP1 but is also found in the cytoplasm, probably due to its affinity for RNA. Consistently, its interactome includes numerous RNA-associated proteins. *Pf*GBP2 is evidently a multifunctional DNA/RNA-binding factor in *Plasmodium*.

## INTRODUCTION

Human malaria, caused by protozoan *Plasmodium* parasites, is responsible for widespread morbidity and almost half a million deaths each year (WHO, 2020). *Plasmodium* lies in an early-diverging lineage which differs greatly from model eukaryotic organisms: it is an obligate intracellular parasite that lives inside host cells for much of its lifecycle, and divides primarily by schizogony rather than conventional binary fission.

*Plasmodium* maintains its genome in conventional linear chromosomes, capped by telomeres that consist of a simple guanine-rich repeat (Figueiredo et al., 2000). These telomeres must be constantly maintained to prevent their degradation during the many replicative rounds of the parasite’s lifecycle. However, *Plasmodium* lacks discernible homologues of almost all of the telomere-binding factors previously identified in model organisms (Zakian, 2012), which control telomere maintenance, recruit or suppress telomerase, enforce transcriptional silencing of adjacent genes via the ‘telomere position effect’ (Gottschling et al., 1990) and suppress the recombination or fusion of DNA ends. In *Plasmodium* the telomere repeat sequence differs slightly from that of the human host (GGGTT(T/C)A instead of GGGTTA), but it is nevertheless likely that specific proteins exist to cap telomeres, monitor their length, regulate their maintenance and mediate their nuclear clustering and tethering, since all these canonical features of telomere biology appear in *Plasmodium* (Bottius et al., 1998; Freitas-Junior et al., 2000).

The first telomeric protein characterized in *Plasmodium* was telomerase itself (Bottius et al., 1998; Figueiredo et al., 2005) and two new proteins were discovered more recently: a zinc-finger protein *Pf*TRZ (Bertschi et al., 2017) and an ApiAP2 transcription factor *Pf*AP2Tel (Sierra-Miranda et al., 2017). Both are particular to *Plasmodium* telomeres, emphasizing the unusual nature of the *Plasmodium* telosome. Identifying additional telomere-binding factors in *Plasmodium* could improve our understanding of telomere biology beyond model organisms.

Importantly, studying *Plasmodium* telomeres could also reveal potential new drug targets, since all single-celled eukaryotes must maintain their telomeres in order to survive. Accordingly, various telomere-targeting drugs that were designed as anti-cancer agents have also been tested against *Plasmodium* (De Cian et al., 2008; Harris et al., 2018). These drugs are frequently designed to target a particular DNA structure called the guanine-quadruplex (G4), which can form in single-stranded guanine-rich sequences such as telomere repeats. G4s occur at eukaryotic telomeres and play important roles in telomere maintenance – hence their potential as anti-cancer targets (Murat and Balasubramanian, 2014). We have reported that a G4-binding drug called quarfloxin kills *Plasmodium* parasites rapidly and potently *in vitro* (Harris et al., 2018), raising the possibility of repurposing it and/or other such drugs as anti-malarials.

Here, we aimed to identify and characterize novel telomere-binding proteins in *Plasmodium falciparum*, using the agnostic approach of pulling down fragments of telomeric chromatin and identifying the associated proteins by mass spectrometry. This method, called Proteomics of Isolated Chromatin fragments, or ‘PICh’, previously identified more than 80 telomere-binding components in human cells (Dejardin and Kingston, 2009). It was adapted to *P. falciparum* – a method that may prove useful in future for identifying other chromatin-domain-specific proteins – and it identified the protein *Pf*GBP2 (PF3D7_1006800). *Pf*GBP2 is an RRM-domain protein whose yeast homolog, ‘G-strand Binding Protein 2’, is known to bind to single-stranded telomeric DNA in *S. cerevisiae* (Lin and Zakian, 1994), as well as binding to mRNAs for nuclear/cytoplasmic shuttling (Windgassen and Krebber, 2003). We confirmed the interaction of *Pf*GBP2 with *Plasmodium* telomere repeats and also with G-rich RNAs *in vitro*. Consistent with this, tagged *Pf*GBP2 was found *in vivo* in the nucleus as well as the cytoplasm of blood-stage *P. falciparum* parasites and interacted with numerous RNA-associated proteins, as well as some DNA-associated proteins. Thus, it seems likely that *Pf*GBP2 plays a role in telomere maintenance, via its binding to telomeric G4s, and also in RNA dynamics.

## RESULTS

### A PICh protocol for *Plasmodium* parasites

Relatively little is known about how *Plasmodium* telomeres are replicated and maintained, making the identification of novel telosome components in *Plasmodium* a priority. Telomere length appears to be a complex trait: there is striking variation in the average length at which telomeres are maintained in different strains of *P. falciparum*, yet their length is relatively stable per strain during *in vitro* culture (Fig. 1a, (Merrick et al., 2012)). This suggests that the trait has a ‘set-point’ – perhaps enforced when new *Plasmodium* strains are generated sexually within a mosquito – and both genetic and epigenetic factors may contribute to telomere maintenance. To investigate the proteins involved in this phenomenon, we set out to identify new telosome components in the *P. falciparum* parasite.

**Figure 1:**
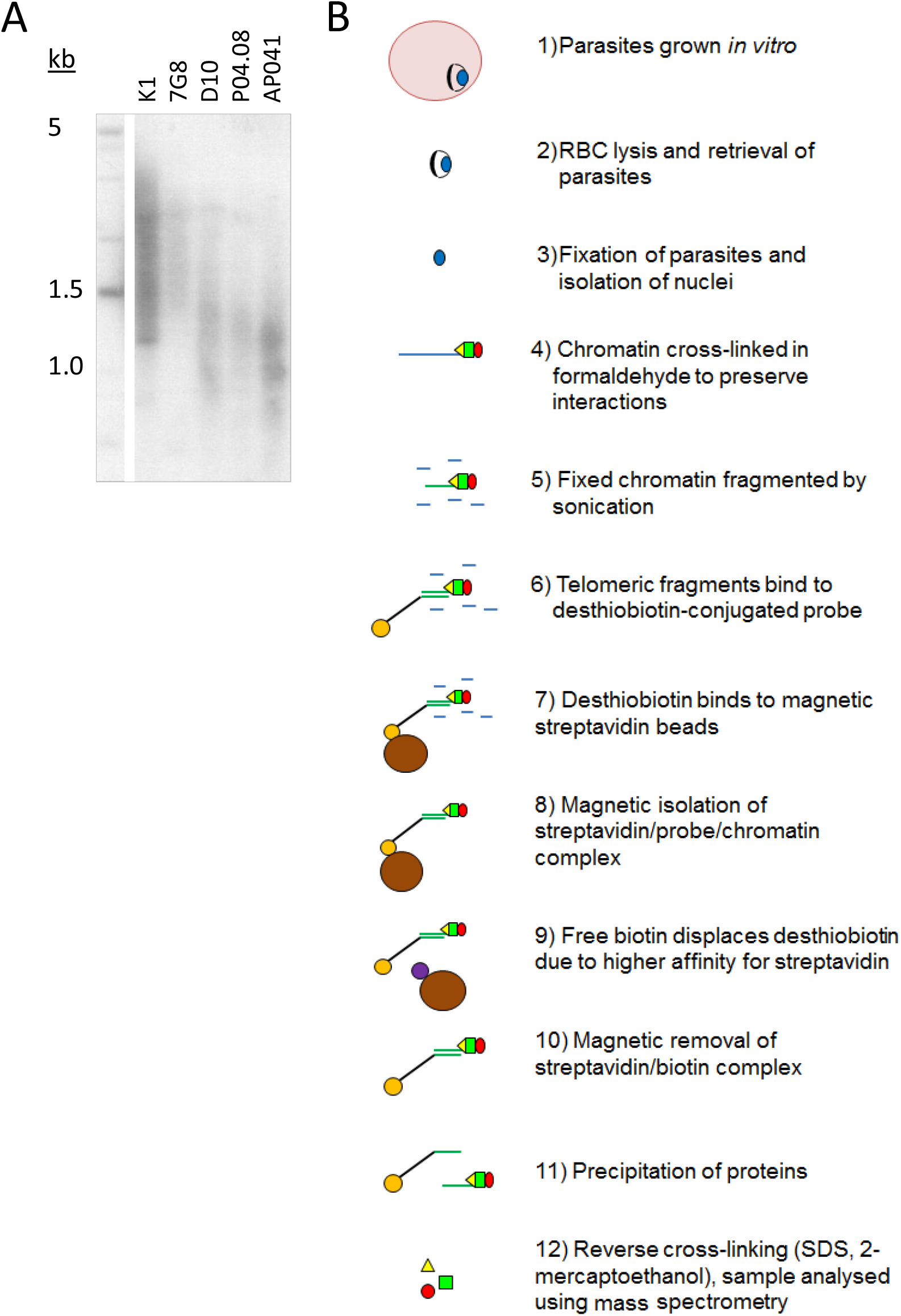
*Plasmodium* telomeres vary in their set-point lengths. (A) Telomere Restriction Fragment Southern blot showing variation in telomere lengths in geographically diverse strains of *P. falciparum* (K1, Thailand; 7G8, Brazil; D10, Papua New Guinea; P04.08, Senegal; AP041, Nigeria). (B) Schematic showing the process of PICh in *P. falciparum*.

The published protocol for PICh in Hela cells was adapted for *P. falciparum* (Fig. 1b), using DNA probes adapted to the *Plasmodium* telomere sequence (GGGTT(T/C)A with 67% T, 33% C at the variable position), and crosslinking the chromatin after releasing parasites from host erythrocytes and washing then thoroughly to reduce contamination from host haemoglobin. Parasite chromatin extracts were probed in parallel with either a telomere-repeat probe or a scrambled probe and the proteins thus purified were identified by mass spectrometry. Yields were initially very limited (a first experiment produced only five *P. falciparum* proteins, including histones and other highly-abundant proteins like elongation factor 1 alpha, which were largely similar in the telomere-probe and control-probe conditions). However, a second experiment using the alternative method of gel-aided rather than filter-aided sample preparation for mass spectrometry gave a much greater yield of over 30 *P. falciparum* proteins. There remained a high representation of histones and other abundant proteins (Table S1), and indeed similar issues were reported when *Pf*TRZ and *Pf*AP2Tel were previously identified via a different methodology (pull-down from nuclear extract onto telomeric versus scrambled DNA probes). In these studies only 12 out of 109 (Bertschi et al., 2017) or 7 out of 100 (Sierra-Miranda et al., 2017) of the proteins identified were telomere-probe-specific, but *bona fide* telomere proteins could nevertheless be selected. Similarly, one interesting candidate protein emerged from the PICh dataset.

### PICh identifies *Pf*GBP2 as a putative telomere-binding protein

The most promising candidate protein found by PICh was encoded by the gene PF3D7_1006800: a putative homologue of *S. cerevisiae* GBP2. *Pf*GBP2 is a protein of 246 amino acids encoding two RNA Recognition Motif (RRM) domains. These domains are well-characterized to occur in proteins that bind to single-stranded nucleic acids, either DNA or RNA (Query et al., 1989). The RRM structure consists of two helices and four strands in an alpha/beta sandwich which can bind to a strand of nucleic acid, and indeed *Pf*GBP2 was modelled with two RRM domains, joined by a less structured linker region (Fig. 2a). In contrast, *Sc*GBP2 is a larger protein with three RRM domains, the third of which is divergent and acts instead as a protein-protein interaction domain (Martinez-Lumbreras et al., 2016) (Fig. 2b). This third domain is lacking in the *P. falciparum* homolog and both RRM domains in *Pf*GBP2 are actually most homologous to RRM2 in *Sc*GBP2, which is the principal nucleic-acid-binding domain (Martinez-Lumbreras et al., 2016) (Fig. 2c).

**Figure 2:**
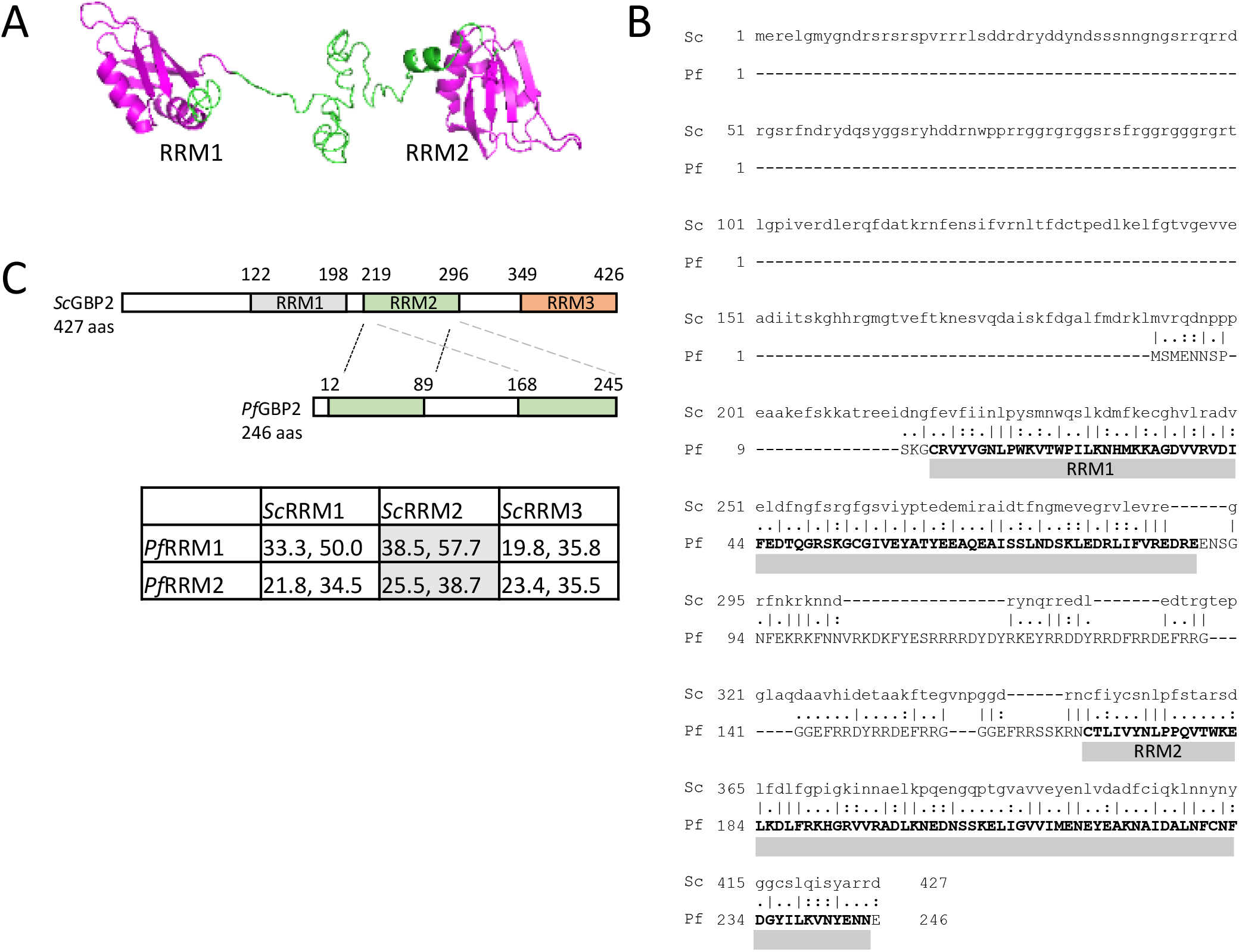
PiCh identifies *Pf*GBP2, a RRM-motif protein. (A) Protein structure model for *Pf*GBP2, modelled using iTASSER (C-score -2.44). (B) Amino acid alignment of *Pf*GBP2 with *Sc*GBP2. Grey bars denotes the regions containing Prosite RRM motifs. (C) Schematic showing the domain structure of *Sc*GBP2 and *Pf*GBP2. Table shows amino acid identity and similarity scores from pairwise alignments of the individual RRM domains: grey highlighted boxes show that both RRM domains from *Pf*GBP2 score most highly against *Sc*GBP2 RRM2.

Several transcriptomic datasets collated in PlasmoDB (Aurrecoechea et al., 2009) show that *PfGBP2* is expressed at all lifecycle stages, while polysomal RNA studies report that the gene transcript is maximally translated in trophozoites (Painter et al., 2018). In proteomic studies, *Pf*GBP2 is in the nuclear proteome, as expected (Oehring et al., 2012). Overall, data from multiple sources including protein modelling, transcriptomics and proteomics all supported the probability that *Pf*GBP2, being nuclear, nucleic-acid-binding and maximally expressed at replicative stages, could be a *bona fide* telomere protein.

### Recombinant *Pf*GBP2 binds to G-rich telomere sequences

To confirm that *Pf*GBP2 can actually bind to telomeric DNA, we produced a recombinant version of the protein (Fig. 3a). Histidine-tagged *Pf*GBP2, expressed in *E. coli*, could be purified primarily as a full-length protein of ∼35 kDa (predicted MW of 34 kDa including tags; some breakdown products were also co-purified, probably as single RRM domains after degradation at the flexible region). Extracts containing *Pf*GBP2 were then used in electrophoretic mobility shift assays (EMSAs) on a DNA oligonucleotide consisting of a series of G-rich telomere repeats. This DNA was clearly retarded due to protein binding, which was not the case with either a scrambled oligonucleotide or a sequence comprised of A and T bases only (Fig. 3b). Thus, *Pf*GBP2 evidently has a tropism for G-rich DNA, and furthermore for G-triad motifs (e.g. GGGTTTA), since scrambling this sequence abrogated binding.

**Figure 3:**
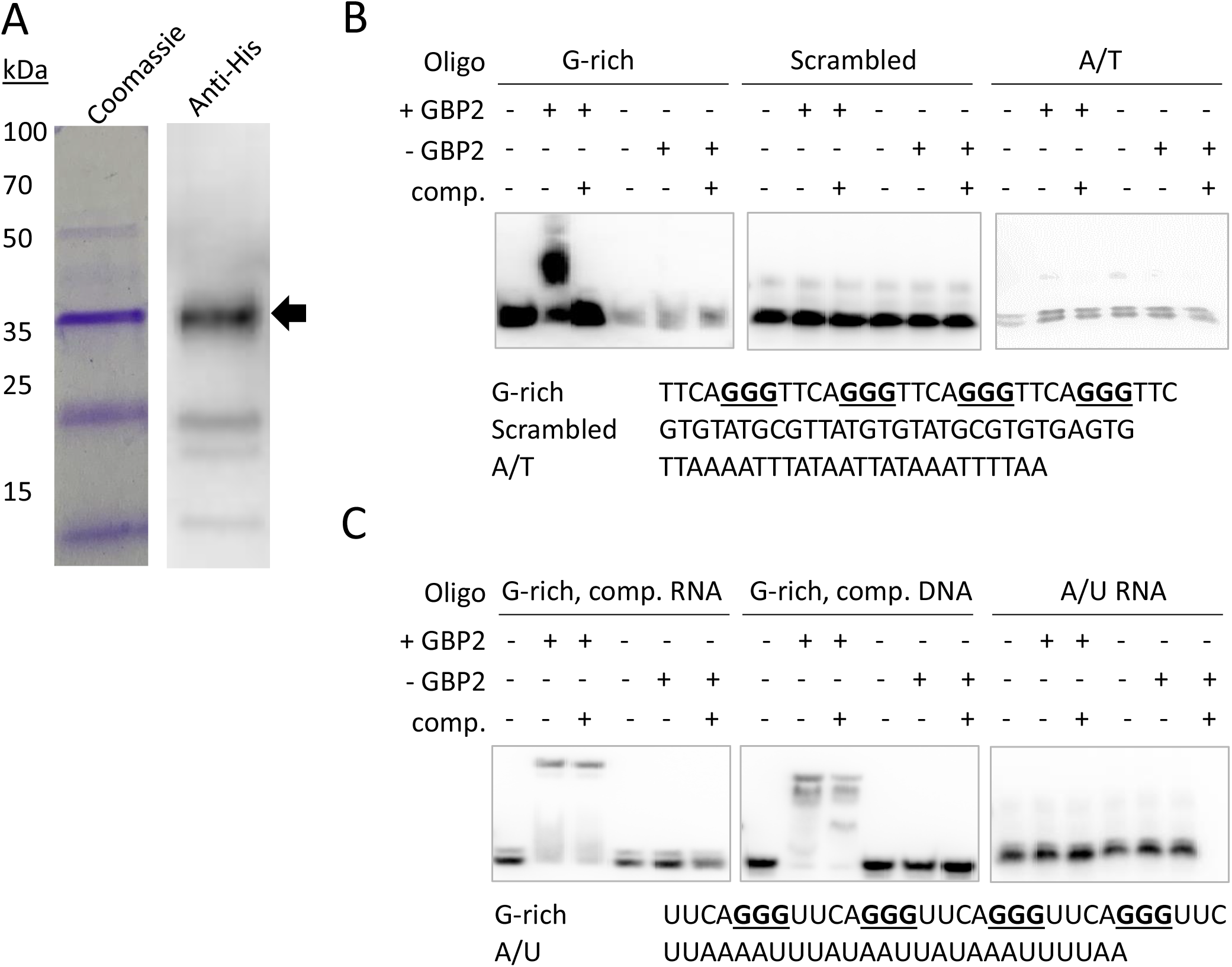
*Pf*GBP2 binds to telomeric DNA and RNA sequences. (A) Recombinant 6x His-tagged *Pf*GBP2 (full-length protein marked with arrow), expressed in *E. coli* and purified via nickel resin. Coomassie-blue-stained gel and western blot against the 6x His-tag. Images are representative of several independent preparations. (B) EMSA assays with the indicated oligonucleotides and bacterial extract containing *Pf*GBP2, or equivalent extract containing no recombinant protein in the control condition. ‘Comp’, unlabeled competitor DNA. Images are representative of several independent experiments. (C) EMSA assay as in (B), using RNA instead of DNA. Competition was attempted with an excess of either unlabeled RNA or unlabeled DNA.

RRM-domain proteins commonly bind RNA as well as DNA, so we investigated whether *Pf*GBP2 might also bind to RNA: EMSAs performed with G-rich telomere repeat RNA oligos showed that this was indeed the case (Fig. 3c). Unlike the behavior seen in the DNA EMSA, *Pf*GBP2 was not efficiently competed off by unlabeled RNA, and was only partially competed off by unlabeled DNA.

### Recombinant *Pf*GBP2 binds to G-quadruplex DNA

Next, we sought to determine whether the G-rich telomere repeat sequence was actually folded into a G4 when bound to *Pf*GBP2, since it was theoretically possible that the DNA would be bound either as a G4 or as a linear strand. Two independent assays showed that the *Plasmodium* telomere repeat sequences used here can indeed fold into G4s in the presence of K^+^ ions, which are required to stabilize quadruplex structures. Figure 4a shows a dot-blot with the G4-structure-specific antibody BG4 (Biffi et al., 2013), while figure 4b shows fluorescent emission from a G4-specific dye, thioflavin T, which induces G4 folding and fluoresces strongly only when bound to a G4 (Mohanty et al., 2013; Renaud de la Faverie et al., 2014). In both these assays, two variants on the *Plasmodium* telomere repeat (GGGTT(T/C)A) were tested, with different representations at the variable T/C position (‘G-rich 1’ and ‘G-rich 2’, all oligonucleotides are listed in Table S2). Both variants behaved identically: when folded in the presence of K^+^ they showed strong binding to the G4-specific antibody and strong emission from thioflavin T. By contrast, the equivalent treatment in the presence of Li^+^ ions, which destabilize G4s, yielded lower signals in both assays, similar to those of a control A/T-only sequence. We also confirmed that four G-triads were required to form a G4, because the same sequence truncated to just three repeats did not give a strong G4 signal in either assay.

**Figure 4:**
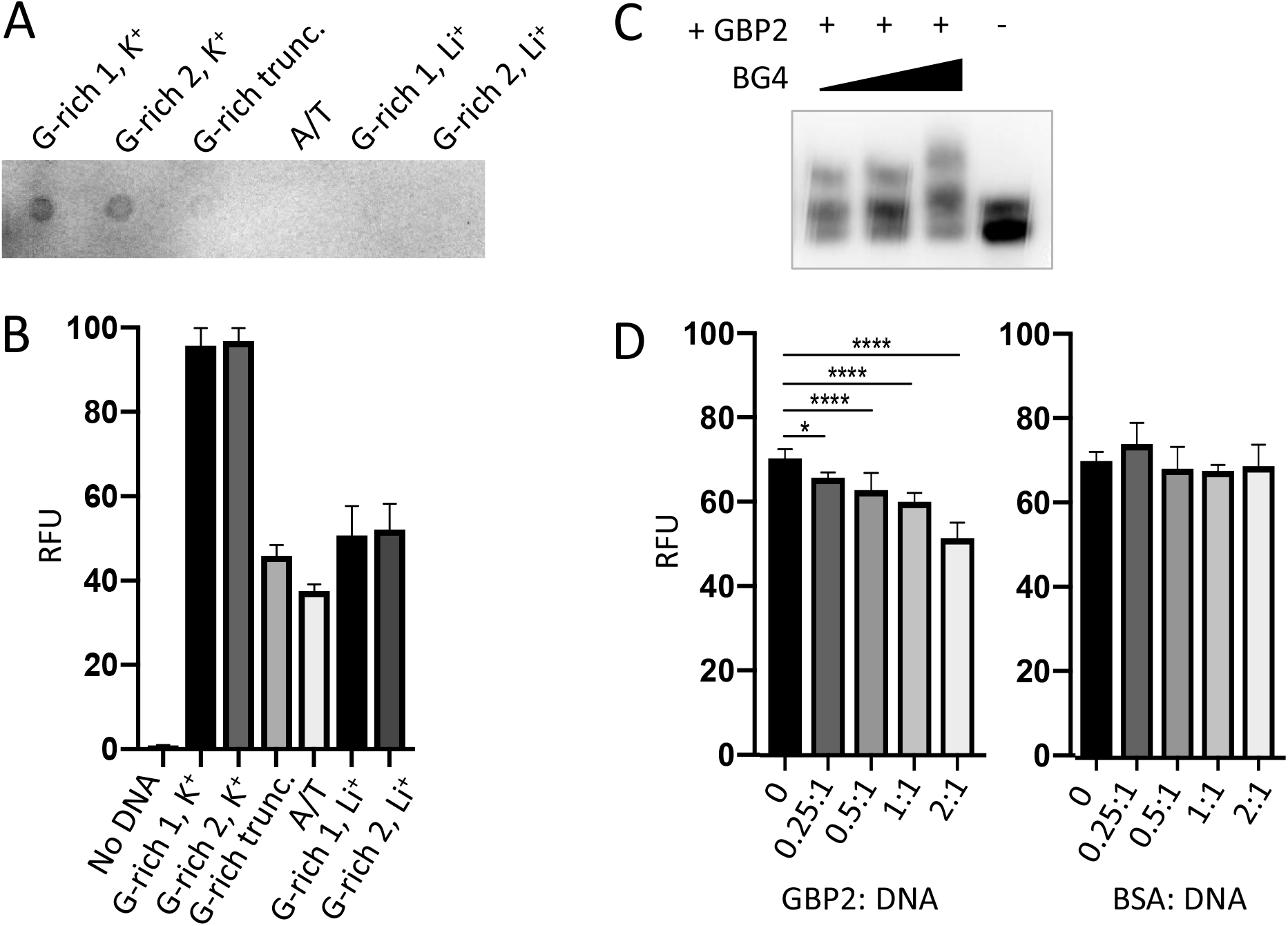
*Pf*GBP2 binds to G4-folded DNA. (A) Dot-blot of the indicated oligonucleotides probed with the G4-specific antibody BG4. Image is representative of triplicate experiments. (B) Fluorescence emission from the indicated oligonucleotides in the presence of the G4-specific dye thioflavin T (ThT). Error bars represent SD from technical triplicates. (C) EMSA assay as in Figure 3B, with BG4 antibody added to the DNA/*Pf*GBP2 complex at 0.5:1, 1:1 and 2:1 molar ratio of antibody to purified *Pf*GBP2. (D) Fluorescence emission from G-rich oligonucleotide 1 bound to thioflavin T, as in (B), with the addition of increasing quantities of purified *Pf*GBP2 or the control protein BSA. Protein:DNA molar ratios between 0.25:1 and 2:1 were tested.

Having confirmed the specificities of these two assays for G4s, the BG4 antibody was then added to the DNA EMSA, where it exerted an additional shift upon the oligo-*Pf*GBP2 complex, showing that the complex indeed contained G4 DNA (Fig. 4c). Finally, *Pf*GBP2 was also able to interfere with thioflavin T emission when added to a mixture of thioflavin T and DNA (Fig. 4d), whereas an irrelevant protein (bovine serum albumin) could not. This interference could potentially occur via *Pf*GBP2 binding to the DNA and dampening the emission from the dye in its G4-bound form; alternatively, it could occur because *Pf*GBP2 actually competes the dye off the G4 motif. In summary, multiple independent assays showed that *Pf*GBP2 is a *bona fide* G4-binding protein.

### *Pf*GBP2 is found in both the nucleus and cytoplasm in erythrocytic parasites

Having characterized *Pf*GBP2 *in silico* and *in vitro*, we proceeded to investigate its properties *in vivo*. A gene knockout of *PfGBP2* was not attempted because this was found to be very deleterious in a recent forward-genetics screen for essential genes in *P. falciparum*, (Zhang et al., 2018): *PfGBP2* mutants had a fitness score of -2.5, only slightly higher than -3 in telomerase reverse transcriptase (TERT) knockouts. Instead, overexpression of the *PfGBP2* gene was attempted in 3D7 parasites, via a tagged version of the gene transfected in episomally in addition to the endogenous copy. No transgenic parasites were obtained after three separate transfections with two different plasmids, carrying *PfGBP2* with two different C-terminal tags (HA and Ty) and two different selectable markers: this strongly suggested that overexpression of tagged *Pf*GBP2 protein was also deleterious. Ultimately, in order to localize the *Pf*GBP2 protein in blood-stage parasites, the endogenous gene was C-terminally tagged with a triple HA tag using the selection-linked integration system (Birnbaum et al., 2017). Correct tagging was confirmed by PCR and the tagged protein was detected in parasites by both western blot and immunofluorescence (Fig. 5).

**Figure 5:**
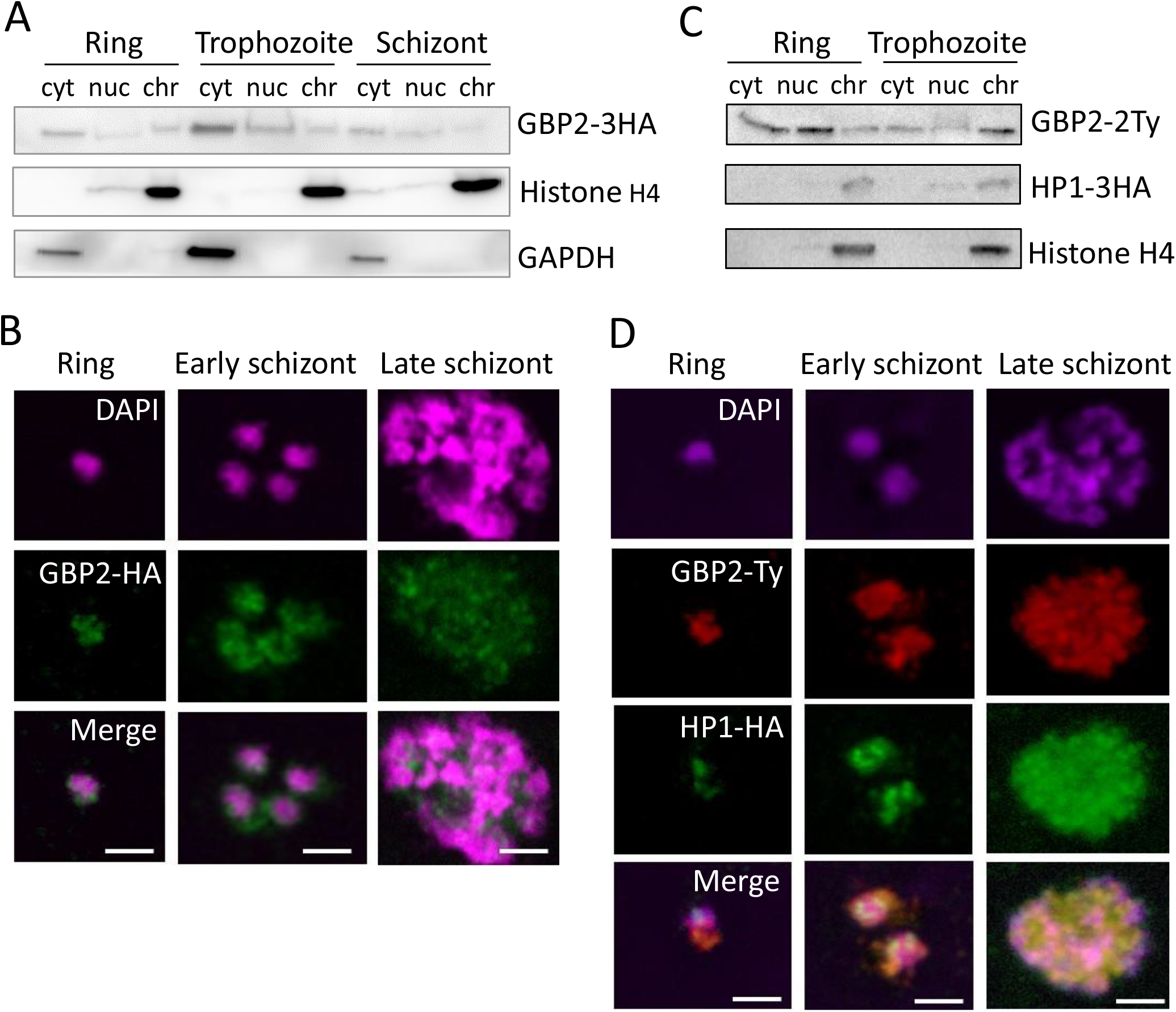
*Pf*GBP2 is found in both the nucleus and cytoplasm in erythrocytic parasites. (A) Western blot of protein fractions (cyt, cytoplasm; nuc, nucleoplasm; chr, chromatin-bound) from ring, trophozoite and schizont 3D7 parasites expressing *Pf*GBP2-3HA. Parallel control blots show histone H4 (nuclear) and glyceraldehyde 3-phosphate dehydrogenase (GAPDH, cytoplasmic). Images are representative of several independent fractionation experiments. (B) Representative immunofluorescence images of ring, trophozoite and schizont 3D7 parasites expressing *Pf*GBP2-3HA, stained with an antibody against the HA tag and DAPI to identify parasite nuclei. Scale bar, 2μm. (C) Western blots as in (A): 3D7 parasites expressing *Pf*GBP2-2Ty and HP1-3HA. (D) Representative immunofluorescence images as in (B), parasites expressing *Pf*GBP2-2Ty and HP1-3HA.

Western blotting revealed *Pf*GBP2-3HA in the nucleoplasm and chromatin-bound fractions of all erythrocytic parasite stages (Fig. 5a), as would be expected for a telomere-binding protein, but it was also found in the cytoplasm at all stages, most prominently in trophozoites. Consistently, *Pf*GBP2-3HA was detected by immunofluorescence in individual parasites as peri-nuclear foci which are characteristic of telomeric factors (Fig. 5b): these appeared at all stages but *Pf*GBP2-3HA was always present in the parasite cytoplasm as well.

To further confirm that the peri-nuclear foci of *Pf*GBP2-3HA did represent telomeres, the *PfGBP2* gene was Ty-tagged in a line already expressing the well-characterised telomeric factor heterochromatin protein 1 (HP1) with an HA tag (Flueck et al., 2009) (Fig. 5c). The two tags, HP1-HA and *Pf*GBP2-Ty, partially colocalised, particularly in late schizonts, with each merozoite bearing a perinuclear focus of both GBP2 and HP1. At earlier stages, however, HP1 foci were discrete, whereas *Pf*GBP2 was dispersed throughout the parasite (Fig. 5d). This was consistent with the fractionation of these parasites showing that HP1 was entirely restricted to the nucleus whereas *Pf*GBP2 was not (Fig. 5c). The tropism of GBP2 for RNA as well as DNA may explain the widespread localization of this protein.

Finally, to define the binding sites of *Pf*GBP2 throughout the genome, chromatin immunoprecipitation (ChIP) was attempted. A ChIP/dot-blot suggested that *Pf*GBP2-3HA was modestly enriched on telomeric DNA (Fig. S1a), but ChIP-seq for either *Pf*GBP2-3HA or *Pf*GBP2-Ty failed to give signals significantly above background at any locus. This compared with strong signals from the established sub-telomeric protein HP1 (Flueck et al., 2009) that was co-expressed in the *Pf*GBP2-Ty line. In a series of gene-directed ChIP experiments (Fig. S1b), HP1 was enriched by over 50-fold at all sub-telomeric loci compared to chromosome-internal loci, whereas *Pf*GBP2 was enriched by only ∼2-fold at sub-telomeric and G4-encoding loci compared to chromosome-internal loci. This demonstrated that the ChIP experiment was conducted correctly but that *Pf*GBP2 did not, in our hands, give a strong enough signal for a meaningful ChIP-seq experiment.

### The interactome of *Pf*GBP2 suggests roles in both DNA and RNA metabolism

To learn more about the potential biological roles of *Pf*GBP2, the HA-tagged protein was immunoprecipitated (IP) and its interactome was obtained via mass spectrometry. Duplicate IP experiments were conducted, yielding a total of 29 reproducible hits specific to *Pf*GBP2 (i.e. absent from an identical control experiment using wildtype parasites) (Fig. 6A, Table S3). A larger group of 187 proteins appeared uniquely in just one of the two *Pf*GBP2 IP experiments (Fig. 6B, Table S3).

**Figure 6:**
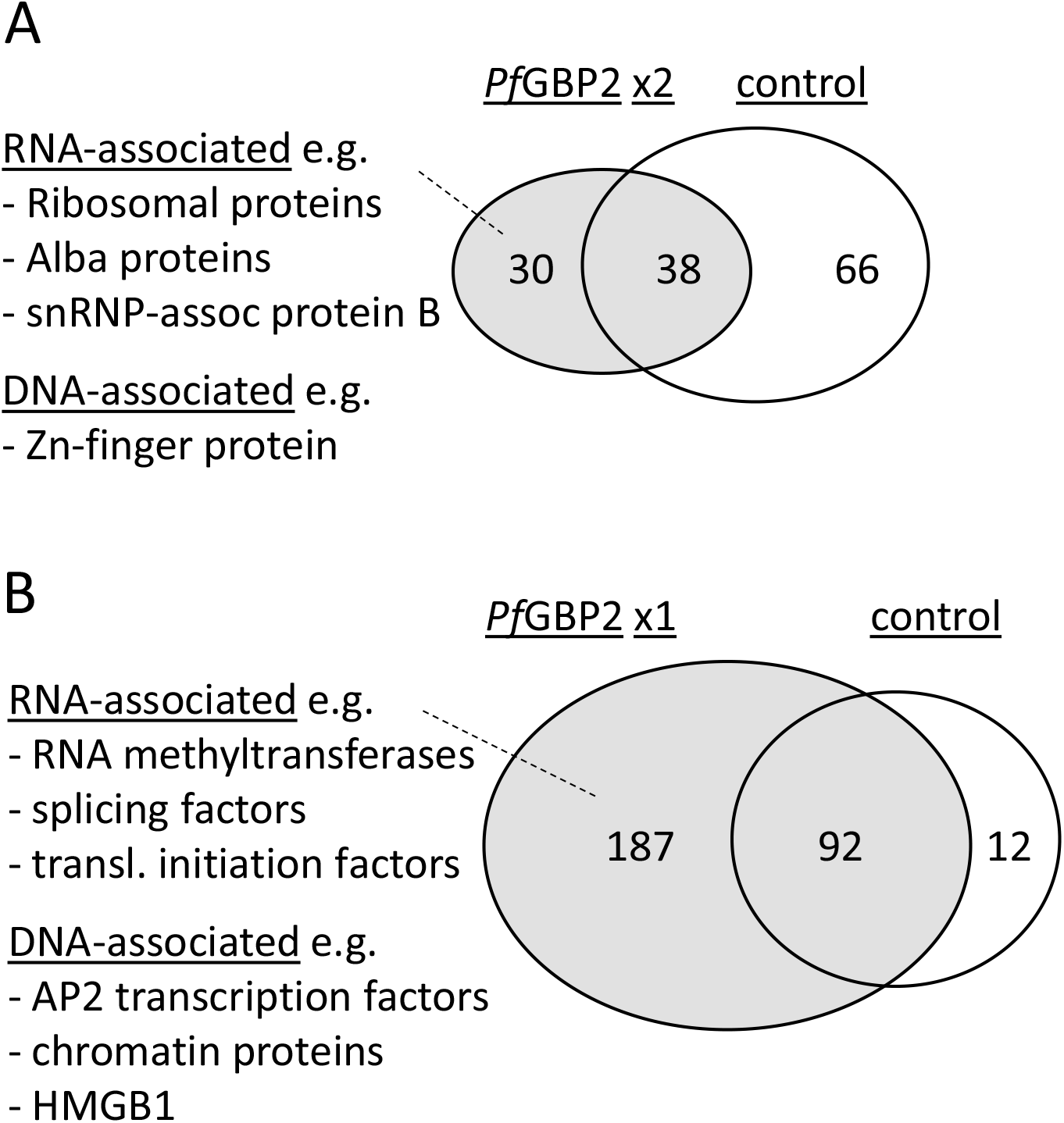
*Pf*GBP2 interacts primarily with RNA-associated proteins. (A) Venn diagram showing the proportion of *Pf*GBP2-interacting proteins found reproducibly in duplicate experiments but absent from the control experiment, with examples of representative proteins. (B) Venn diagram showing the larger number of *Pf*GBP2-interacting proteins found in only one duplicate experiment, with examples of representative proteins.

Amongst the reproducible hits there was a clear preponderance of RNA-associated proteins. Gene ontology terms including ‘cytosolic ribosome’, ‘ribonucleoprotein complex’, and various terms concerning mRNA editing and base modification were enriched in the interactome (Fig. 6A, Table S4). A few DNA-binding proteins were also represented, including a zinc-finger protein (PF3D7_1317400), but DNA-related GO terms were not strongly enriched overall, and the known telomeric proteins *Pf*TRZ or *Pf*AP2Tel did not appear. A broader analysis of all 187 putative *Pf*GBP2-interacting proteins yielded similar results, i.e. a clear enrichment of RNA-associated proteins (Table S4), as well as a few DNA-associated proteins.

These results were compared with those of a recent study that used machine learning to infer a proteome-wide interactome for *P. falciparum* (Hillier et al., 2019). This reported that at least 17 of the top 50 interactors for *Pf*GBP2 were RNA-associated proteins, including several initiation factors and snRNP-associated proteins, while 5 out of 50 were DNA-associated proteins, including a DNA helicase, a DNA repair protein, a transcription factor and the High Mobility Group protein HMGB1. Only 5 out of these 50 top interactors appeared as *Pf*GBP2 interactors in one of our two datasets, including the transcription factor (PF3D7_1426100) and *Pf*HMGB1 (PF3D7_1202900). The latter protein is particularly interesting because in human cells, it was recently reported to interact with telomeric G4 DNA (Amato et al., 2019), raising the possibility that *Pf*GBP2 and *Pf*HMGB1 might cooperate at telomeric G4s. Overall, the interactome strongly suggests that *Pf*GBP2 is present in RNA-binding as well as DNA-binding complexes.

## DISCUSSION

This work set out to identify novel *Plasmodium* telosome components, and subsequently to characterise the GBP2 protein in *P. falciparum*. This involved the development of a ‘PICh’ method to pull down sequence-specific chromatin fragments from *P. falciparum*: a method that may have applications in future studies. PICh did identify a new telosome component, but it did not identify telomerase or other *Plasmodium*-specific telosome proteins, *Pf*TRZ (Bertschi et al., 2017) or *Pf*AP2Tel (Sierra-Miranda et al., 2017), which were both discovered via DNA-mediated pulldowns from parasite extracts. Those two reports did not identify one another’s proteins either, suggesting that no method is entirely comprehensive and that more proteins may be undiscovered. In PiCh, however, the proteins are identified directly from native chromatin rather than from protein extracts that were subsequently re-bound to DNA probes, so there is potential to identify different sets of proteins. In particular, *Pf*GBP2 evidently targets the G-rich telomeric overhang, whereas *Pf*TRZ and *Pf*AP2Tel (Myb- and AP2-domain proteins) bind to double-stranded DNA. The PiCh technique may thus be better-placed to detect components of native telomeric chromatin that are not dsDNA-binders and are not pulled down by conventional DNA probes. Of note, however, a second study published during the preparation of this manuscript did identify *Pf*GBP2 via pulldown from parasite extracts, using a G-quadruplex-forming DNA sequence as the probe (Gurung et al., 2020).

Unlike *Pf*TRZ and *Pf*AP2Tel, *Pf*GBP2 is not unique to *Plasmodium*: homologues exist in eukaryotes including plants, yeast and humans, as well as other apicomplexans. In apicomplexans, GBP2 takes a short form with just two DNA-binding RRM domains. This is also the form found in plants, whereas in *S. cerevisiae* there is a third, divergent RRM domain which mediates protein-protein interaction with the THO/TREX mRNA export complex (Martinez-Lumbreras et al., 2016), and *Sc*GBP2 accordingly has dual functions in telomere binding and mRNA metabolism. Dual roles for such proteins are not unusual: some hnRNPs also bind to both G-rich RNA and telomeric ssDNA, and play roles in both RNA metabolism and telomere stabilisation (Tanaka et al., 2007). Indeed, we present here the first evidence that *Pf*GBP2 binds to G-rich RNA as well as DNA, and we also suggest that *Pf*GBP2 overexpression may be lethal, as *Sc*GBP2 overexpression is also lethal, owing to deregulated mRNA export (Windgassen and Krebber, 2003). Nevertheless, the mRNA shuttling role played by *Sc*GBP2 is probably not directly conserved in *P. falciparum*, since *Sc*GBP2 requires its third domain for recruitment to nascent mRNA via TREX (Hurt et al., 2004), and not all components of the THO/TREX complex have even been identified in *Plasmodium* (Tuteja and Mehta, 2010). Therefore, any interaction with RNA may be mediated differently in parasites.

By contrast, it is clear that the role in telomeric DNA binding *is* conserved among yeasts, plants and apicomplexans. In *S. cerevisiae*, GBP2 lacks an essential telomeric function: it does protect telomeric ssDNA (Pang et al., 2003) but telomeres can still be maintained in its absence, albeit with mislocalisation of the Rap1 protein (Konkel et al., 1995). On the contrary, in plants, the telomere-binding role of GBP2 is essential. In *Nicotiana tabacum*, its loss causes severe developmental and chromosomal abnormalities with defective telomeres (Lee and Kim, 2010). *Pf*GBP2 is more closely homologous to the plant version than the yeast version, sharing 46% similarity with *Nt*GBP2, and the *PfGBP2* gene was essential or near-essential in a *P. falciparum* genome-wide screen (Zhang et al., 2018). However, in their recent report on *Pf*GBP2, Gurung and co-workers were able to achieve a knockout which surprisingly had no growth defect, nor was telomere maintenance affected (Gurung et al., 2020). A viable *P. berghei* GBP2 knockout has also been reported and although its telomeres were not assessed, this parasite line did grow slowly (Niikura et al., 2020).

All these data call into question the expectation that GBP2 might be essential in *Plasmodium* and might play a role in telomere maintenance. However, the knockouts reportedly achieved in both *P. falciparum* and *P. berghei* may have been non-homogenous, since the genetic status of the knockout populations was not confirmed after long-term growth. A salient example in the literature reports the knockout of another essential telomeric protein, telomerase, via disruption of the *TERT* gene in *P. berghei*. Knockouts were briefly detected in bulk culture, but could never be cloned out before they were outgrown by healthier non-knockout parasites (Religa et al., 2014). This was probably because the telomeres in the knockout parasites quickly became critically degraded, so the authors concluded that *Pb*TERT was essential. It would be interesting to establish whether outgrowth of non-knockout parasites could also occur if *GBP2* knockout parasites are debilitated by telomere loss.

Whether or not the telomere-binding role of *Pf*GBP2 is essential, the role clearly exists. On this point our work is consistent with that of Gurung *et al*., and also with a 2015 study (published in Spanish and not indexed via PubMed) which previously identified *Pf*GBP2 *in silico* as a putative telosome component and confirmed that it binds specifically to G-rich telomere-repeat oligos *in vitro* (Calvo and Wasserman, 2015). The same property has been tested in other apicomplexans as well: *Eimeria* GBP2 was found at telomeres (via semi-quantitative ChIP-PCR (Zhao et al., 2014)), while *Cryptosporidium* GBP2 bound to telomeric DNA *in vitro* and specifically required its first RRM domain to do so (Liu et al., 2009). Our work goes further in examining the quadruplex-binding capacity of *Pf*GBP2: we conducted two independent assays to detect folded G4s in *Pf*GBP2-DNA complexes. The exact G4 binding mode of the protein is unknown, but if *Pf*GBP2 can directly compete with ThT to bind G4s (which is one explanation for the data in figure 4d), then this would suggest an end-stacking mode, because thioflavin T is thought to end-stack onto the terminal G-quartet of a G4 (Mohanty et al., 2013). Further biophysical studies would be needed to confirm this. Finally, our work also goes further in exploring the binding of *Pf*GBP2 to RNA as well as DNA G4s. An affinity for RNA explains the broad cellular location of this protein, and is consistent with the presence of many RNA-binding proteins in the *Pf*GBP2 interactome,

The biological implications of *Pf*GBP2’s clear affinity for DNA/RNA G4s still warrant further study. Gurung *et al*. reported that the G4 affinity is not restricted to telomeres: the protein was initially identified via pulldown on a non-telomeric G4, and it was then found throughout the genome via ChIP-seq (Gurung et al., 2020), although surprisingly the original G4 sequence used in the pulldown did not appear in the ChIP results. These authors reported that *Pf*GBP2 bound very broadly throughout the genome with an extreme enrichment of 500-2000 fold over input: this is an order of magnitude greater than that seen with the *bona fide* sub-telomeric protein HP1 (Flueck et al., 2009). By stark contrast, we were unable to obtain a meaningful ChIP signal, even when *Pf*GBP2 was identically C-terminally tagged in a chromatin preparation from which HP1 could be ChIPed with over 50-fold enrichment. Since Gurung *et al*. did not perform a similar ChIP control, the disparity between these two very similar experiments remains unexplained. Nevertheless, if *Pf*GBP2 does indeed bind very broadly to G-rich sequences throughout the *P. falciparum* genome, the protein could play interesting roles in G4 metabolism beyond telomeres.

Overall, the data presented here, together with the literature on GBP2 proteins across eukaryotes, indicate a triple role for *Pf*GBP2 – in telomeric G4 binding, in pan-genomic G4 binding, and in G4-RNA binding. *Pf*GBP2 is the first G4-binding protein to be identified in *Plasmodium*, and only the third protein, beside telomerase, to be identified as part of the divergent telosome in *Plasmodium*.

## LIMITATIONS OF THE STUDY

The full functional characterisation of this protein awaits further work. A mechanistic role for *Pf*GBP2 in RNA metabolism, such as RNA processing or shuttling, has not been investigated, and nor has its exact role at telomeres (although a complementary paper already reported no effect upon telomere length). In addition, although there is strong evidence that this is a true G4-binding protein, its binding mode has not been evaluated, nor has any preference for particular structures such as parallel vz antiparallel quadruplexes.

## AUTHOR CONTRIBUTIONS

JES – Designed, optimized and conducted PiCh experiments

ALJ – Conducted recombinant protein production, EMSA, ThT fluorescence, dot-blotting, western blotting and co-immunoprecipitation experiments

LER – Cloned the expression vector and conducted recombinant protein production

FIGT – Designed, optimized and conducted ChIP

SRH – Coordinated and analysed data from mass spectrometry on PiCh samples

CJM – Designed the study, conducted experiments (including Southern blotting, cloning, transfection and immunofluorescence assays), analysed data, made figures and wrote the manuscript.

## ACKNOWLEDGEMENTS & FUNDING

We acknowledge the Cambridge Centre for Proteomics and the Liverpool Centre for Proteome Research, particularly Philip Brownridge for expert assistance with PiCh mass spectrometry; Jerome Dejardin for helpful comments and advice regarding PICh; Till Voss (Swiss TPH) for the HP1-HA parasite line; Richard Bartfai and Jonas Gockel (Radboud University) for help with ChIP-seq; Christian Happi (Redeemers’ University) and the group of Dyann Wirth (Harvard University) for supplying parasite genomic DNAs from Nigeria and Senegal.

The work was supported by the UK Medical Research Council [grant MR/L008823/1 to CJM] and UK Biotechnology and Biological Sciences Research Council [grant BB/K009206/1 to CJM].

## DECLARATIONS OF INTEREST

The authors declare no conflict of interest.

## MATERIALS AND METHODS

### Parasite culture and transfection

Laboratory strains of *P. falciparum*, 3D7, HB3, Dd2, K1, 7G8 and D10 were obtained from the MR4 repository (www.beiresources.org). 3D7 was used for all experiments except the telomere Southern blots, which used genomic DNA from other strains. Parasites were maintained *in vitro* in human O+ erythrocytes at 4% haematocrit in RPMI 1640 medium supplemented with 25mM HEPES (Sigma-Aldrich), 0.25% sodium bicarbonate, 50 mg/L hypoxanthine (Sigma-Aldrich), 0.25% Albumax (Invitrogen) and 5% heat-inactivated pooled human serum, using standard procedures (Trager and Jensen, 1976).

Transfections were carried out after synchronization with 5% sorbitol and then maturation to highly synchronous late-stage trophozoites/schizonts. Transgenic parasites were generated by allowing these cultures to invade erythrocytes pre-loaded with 50 - 100 μg plasmid DNA as previously described (Deitsch et al., 2001). Parasites were allowed to grow for 48 hours before being exposed to drug selection, and then maintained with 5nM WR99210 (Jacobus Pharmaceuticals). For pSLI-mediated gene tagging, transfectants were subsequently selected with neomycin, as previously described (Birnbaum et al., 2017), to select parasites carrying the genome-integrated construct. 2µg/ml blasticidin (Invitrogen) was also used to select for simultaneous expression of HP1-3HA in the HP1-3HA+GBP2-2Ty line.

### Telomere restriction fragment Southern blotting

Genomic DNA was extracted from parasites using the QIAamp DNA Blood Mini Kit (Qiagen), digested with restriction enzymes *Alu*I, *Dde*I, *Mbo*II and *Rsa*I, then blotted with a probe specific for telomeres as described previously (Bottius et al., 1998; Figueiredo et al., 2002).

### Proteomics of Isolated Chromatin Segments (PICh)

PICh assays were carried out essentially as described by Dejardin and Kingston (Dejardin and Kingston, 2009), with *Plasmodium*-specific modifications. A full step-by-step PiCh method can be obtained from https://www.epigenesys.eu/images/stories/protocols. Briefly, parasite cultures were expanded and synchronized with two rounds of sorbitol treatment to yield 1L of synchronous late-stage trophozoites at 9% parasitaemia. Parasitized cells were collected by centrifugation and washed in PBS-PMSF, prior to erythrocyte lysis by addition of saponin to 0.1%. Free parasites were then collected by centrifugation and washed four times in PBS-PMSF, before being crosslinked for 30 mins in 3.7% formaldehyde/PBS-PMSF. Thereafter samples were treated as previously described (Dejardin and Kingston, 2009) with the following critical parameters: RNAse incubation: 2 h at room temperature. Sonication: Total “on” time of 15 mins (4 × 7.5min), 30s on, 30s off. Chromatin preparations were split in two (1x target, 1x control) and hybridized with 30μl of probe per sample (a 50-fold molar excess). Probe sequences are provided in Table S2. Probe-chromatin complexes were captured magnetically, washed, eluted and then isolated by TCA precipitation. Protein pellets were de-crosslinked by boiling in 2% SDS, 0.5M 2-mercaptoethanol, 250mM Tris buffer for 30 mins.

### PiCh protein digestion via filter/gel-aided sample preparation and mass spectrometry

De-crosslinked proteins were subjected to either filter-aided sample preparation (FASP) according to the methods of Mann and coworkers (Wisniewski et al., 2009), or gel-aided sample preparation (GASP) following the methods of Fischer and Kessler (Fischer and Kessler, 2015). In the FASP method, samples were processed using a FASP Protein Digestion Kit (Expedeon, Cambridgeshire), following the manufacturer’s procedure. GASP was performed by adding acrylamide 40% (w/v) (Sigma-Aldrich) 1:1 v/v to the sample, enabling formation of protein-containing polyacrylamide plugs upon polymerization using ammonium persulphate and TEMED (Sigma-Aldrich). Gel plugs were then diced by spinning at 14,000 xg through plastic mesh, before being washed using two successive washes with 6 M urea and 100 mM ammonium bicarbonate in 50% acetonitrile, and subjected to in-gel digestion. Peptides extracted from gel pieces were dried under vacuum, dissolved in 0.1 % formic acid and run using a Q-Exactive hybrid mass spectrometer (Thermo Fisher Scientific), coupled online to nanoflow HPLC. For both FASP and GASP-derived peptides, the mass spectrometer was operated in a ‘top10’ mode, whereby the ten most abundant new precursors observed per survey scan are subjected to product ion analysis by collisional dissociation (Michalski et al., 2011). Product ion spectra were then subjected to parsing by Mascot Distiller using standard settings for high resolution product ion spectra as recommended by the manufacturer, and database searching using an in-house Mascot server (Matrix Sciences, London), against a hybrid database comprised of sequences derived from *P. falciparum* (download date 20th July 2015), alongside common contaminant proteins from artefactual sources frequently seen in pulldown proteomics experiments (Mellacheruvu et al., 2013). Data were compared using Scaffold Q+ (v. 4.3.3, Proteome Software, Portland IR).

### Protein modelling

Structural modelling of *Pf*GBP2 was conducted using I-TASSER (Iterative Threading ASSEmbly Refinement) (Yang et al., 2015). Queries were submitted via the online server (http://zhanglab.ccmb.med.umich.edu/I-TASSER/) and modelling was conducted *ab initio* without optional guide templates or specification of secondary structure. Queries were submitted in October 2018.

### Plasmid construction

To clone the *PfGBP2* (*PF3D7_1006800*) gene for recombinant protein production, the full-length transcript minus the stop codon was amplified by PCR from *P. falciparum* cDNA and cloned into the pET-28a+ expression vector between the *Bam*HI and *Xho*I sites, resulting in a construct with dual 6xHis tags at the N and C termini. To clone plasmids for 3’ HA or Ty tagging of the endogenous *PfGBP2* gene via the pSLI system, the latter half of the gene was cloned into a pSLI 3’ HA tagging vector (Birnbaum et al., 2017) between the *No*tI and *Kpn*I sites. Subsequently, the 3’ half of the gene downstream of an endogenous *Bgl*II site, together with the HA tag, were excised and replaced by the same gene portion with a 2xTy tag (this fusion having been previously generated in an episomal overexpression vector which was not tolerated by 3D7 parasites). All primer sequences are provided in Table S2.

### Recombinant protein production

The pET-28a+ expression construct was transferred into BL21(DE3)/pLys strain (Stratagene) and protein production was induced at 37°C with 1 mM IPTG (isopropyl-β-D-thiogalactopyranoside) for 3h. Bacteria were lysed with Bugbuster reagent (Merck Millipore) plus complete protease inhibitors (Roche), and purification was conducted using gravity flow over nickel affinity resin (Thermo-Fisher Scientific) as previously described (North et al., 2005). Purified protein was further concentrated using Amicon Ultra Centrifugal Filter Units (Merck Millipore).

### Western blotting

Parasite fractions for western blotting were prepared as previously described (Voss et al., 2002). Samples were loaded onto 4-12% polyacrylamide gels and electrophoresed at 100V for 60 mins. Electrophoretic transfer to nitrocellulose membrane was carried out at 100V for 60 mins. Membranes were blocked in TBST with 5% milk protein and probed with the following antibodies: 1:2000 anti-Ty1 (Invitrogen), then 1:1500 goat anti-mouse IgG-HRP (Dako); 1:1000 anti-HA (Roche), then 1:1500 goat anti-rat IgG-HRP (Biolegend); anti-histone H4 (Abcam), then 1:1000 goat anti-rabbit IgG-HRP (Abcam); or 1:1000 13.3 anti-GAPDH (European Malaria Reagent Repository), then 1:1500 goat anti-mouse IgG-HRP (Dako). Membranes were washed for 3 × 5 mins in TBST after each antibody step. Clarity Western ECL substrate (Bio-Rad) was added for 3 mins and blots were imaged using a FluorChemM chemiluminescent detection camera (ProteinSimple).

Recombinant protein was blotted with anti-His antibody using the same method: 1:2000 mouse anti-tetra-His IgG (Qiagen); 1:1500 goat anti-mouse IgG-HRP (Dako). Coomassie staining of recombinant protein after gel electrophoresis was performed by addition of 0.1% Brilliant blue R-250 for 20 mins (Fisher), then de-staining in 40% methanol 10% glacial acetic acid.

### Electrophoretic Mobility Shift Assay (EMSA)

EMSAs were optimized and performed using a LightShift optimization and control system (Thermo Scientific). Protein extracts containing *Pf*GBP2, and control extracts lacking the recombinant protein, were made as above. Crude extracts in Bugbuster reagent were purified using HisPur Ni-NTA resin (Thermo Scientific) and run through a Poly-Prep Chromatography Column (BioRad) by gravity. Purified GBP2 protein extract was used for all EMSAs.

Oligonucleotides were labelled using a 3’ biotin end-labelling kit (Thermo Scientific). Binding reactions were carried out at room temperature with 1µg of GBP2 in the presence of 50ng dIdC. Reactions were pre-incubated for 5 mins prior to the addition of 20 fmol of labelled probe, then incubated for a further 20 mins at room temperature. Unlabeled competitor oligonucleotides were added in 200-fold excess relative to probe. Reactions were then run at 100V on a cooled 0.5x TBE-acrylamide gel (4-12% gradient) for 100 mins. Samples were blotted onto nylon membrane (Perkin Elmer) at 380 mA for 60 mins, crosslinked under UV (125mJ) and then blocked, washed and developed using a LightShift chemiluminescent detection kit (Thermo Scientific). EMSA supershift assays were performed similarly, with prior 1h incubation of the biotinylated oligonucleotide with the anti-G4 antibody BG4 (Merck Millipore).

### Dot blotting

To allow G4 folding, DNA oligonucleotides were heated to 90°C for 5 mins before the addition of 100μM Tris buffer pH 7.8 and 100μM KCl, then cooled from 90°C to room temperature at a rate of 5°C/5 min. Alternatively, oligonucleotides were folded in increasing concentrations of LiCl instead of KCl, up to 1M. 5µl of oligonucleotides (1μM) were then spotted on to nitrocellulose membrane (Perkin Elmer) and crosslinked under UV (125mJ) for 5 mins. Membranes were washed and blocked as per western blotting protocol and probed with 1:1500 BG4 (Merck Millipore), 1:1500 DYKDDDK tag (anti-flag, Cell Signalling), and 1:1500 Goat anti-rabbit IgG-HRP (Abcam).

### Thioflavin T fluorescence assay

Oligonucleotides at 20μM were treated with KCl or LiCl as above for dot blotting, then mixed with Thioflavin T (Sigma Aldrich) at a final concentration of 80μM and incubated at room temperature for 5 mins. 40μl of each oligonucleotide mixture was transferred in triplicate to the wells of a 96 well black, Uclear plate (Greiner), and analyzed using a FLUOstar Omega plate reader (BMG Labtech) at Ex. 420nm, Em. 480nm. *Pf*GBP2 competition assays were performed in the same way, with the addition of increasing concentrations of purified *Pf*GBP2, or BSA as a control, prior to the addition of ThT.

### Immunofluorescence

Parasitized erythrocytes were smeared onto microscope slides and fixed in 4% formaldehyde/PBS for 10 mins, rinsed twice in PBS, treated with 0.03% triton/PBS for 10 mins, blocked with 1% BSA/PBS for 30 mins, then incubated with the following antibodies: 1:500 anti-Ty1 (Invitrogen), then 1:1000 Alexa Fluor 546-conjugated anti-rat IgG (Thermo Fisher Scientific); and/or 1:500 anti-HA (Roche), then 1:1000 Alexa Fluor 488-conjugated anti-rat IgG (Thermo Fisher Scientific). Slides were washed for 3 × 5 mins in PBS after each antibody step and in the penultimate wash 2μg/ml 4’,6-diamidino-2-phenylindole (DAPI) (Molecular Probes) was added. Slides were mounted with ProLong Diamond antifade mountant (Thermo Fisher Scientific) and imaged with a Zeiss LSM700 Confocal Microscope.

### ChIP-seq

#### Chromatin preparation

Cultures of 1.6-3.6×10^9^ sorbitol-synchronized parasites at 30-36 hpi were used for ChIP. Chromatin was crosslinked with 1% formaldehyde in culture media for 10 minutes at 37°C, then quenched with glycine at a final concentration of 0.125 M. Parasites were extracted by lysis with 0.05% saponin in PBS. Nuclei were extracted by gentle homogenisation in cell lysis buffer (10mM Tris pH 8.0, 3mM MgCl_2_, 0.2% NP-40, 1x Pierce protease inhibitor (Thermo Fisher)) and centrifugation at 2000 rpm for 10 minutes in 0.25 M sucrose cushion in cell lysis buffer. Harvested nuclei were snap-frozen in 20% glycerol in cell lysis buffer. Thawed nuclei were resuspended in sonication buffer (50mM Tris-HCl, 1% SDS, 10mM EDTA, 1x protease inhibitor (Sigma-Aldrich), pH 8.0) and sonicated for 20-24 cycles of 30s ON, 30s OFF (setting high, BioruptorTM Next Gen, Diagenode) (Fraschka et al., 2018).

#### Chromatin immunoprecipitation

Each ChIP reaction was set up with 500 ng sonicated chromatin incubated in incubation buffer (0.15% SDS, 1% Triton-X100, 150mM NaCl, 1mM EDTA, 0.5mM EGTA, 1x protease inhibitor (Sigma-Aldrich), 20 mM HEPES, pH 7.4) with either 400 ng of α-HA (Roche 12158167001) or 1μl α-Ty (BB2, in-house hybridoma supernatant), together with 10 μL protA and 10 μL protG Dynabeads suspension (Thermo Fisher Scientific). For each sample, eight ChIP reactions were prepared and incubated overnight rotating at 4 °C. Beads were washed twice with wash buffer 1 (0.1% SDS, 0.1% DOC, 1% Triton-X100, 150 mM NaCl, 1 mM EDTA, 0.5 mM EGTA, 20 mM HEPES, pH 7.4), once with wash buffer 2 (0.1% SDS, 0.1% DOC, 1% Triton-X100, 500 mM NaCl, 1 mM EDTA, 0.5 mM EGTA, 20 mM HEPES, pH 7.4), once with wash buffer 3 (250 mM LiCl, 0.5% DOC, 0.5% NP-40, 1 mM EDTA, 0.5 mM EGTA, 20 mM HEPES, pH 7.4) and twice with wash buffer 4 (1 mM EDTA, 0.5 mM EGTA, 20 mM HEPES, pH 7.4). Each wash step was performed for 5 min at 4°C while rotating. Immunoprecipitated chromatin was eluted in elution buffer (1% SDS, 0.1M NaHCO3) at room temperature for 20 min. The eluted chromatin samples and the corresponding input samples (sonicated input chromatin containing 500 ng DNA) were de-crosslinked in 1% SDS / 0.1M NaHCO3 / 1M NaCl at 65°C for at least 4h while shaking, followed by column purification (PCR Purification Kit, Qiagen) and elution in 200ul EB buffer.

#### Quantitative PCR

qPCRs were performed with 5µL ChIP-ed DNA against a 10x dilution series of input DNA using iQ™ SYBR Green Supermix (Biorad) together with primers (Table S2) mixed according to manufacturers’ instructions on a C1000 Touch CFX96 Real-Time System (Biorad).

### Co-immunoprecipitation and mass spectrometry

800ml of 3D7 WT and 3D7 GBP2-3HA cultures were saponin-treated to release the parasites (1-2 × 10^10^ parasites per sample, conducted in biological duplicate for GBP2). Parasites were re-suspended in lysis buffer (1% Triton, 50mM HEPES, 150mM NaCl, 1mM EDTA) and subjected to a freeze-thaw cycle three times, before treating with 1 unit of DNase1 for 10mins at 37°C (Thermo Fisher Scientific). Samples were then centrifuged for 30 mins at 4°C at 14500 rcf. Supernatant was added to Protein G magnetic beads (Pierce) pre-washed three times in wash buffer (0.1% Triton, 50mM HEPES, 150mM NaCl) and incubated for 1 h at 4°C. Magnetic beads were removed by magnet and 1mg/ml of anti-HA antibody (Roche) was added to the proteins for incubation overnight at 4°C. Following incubation, a new aliquot of Protein G magnetic beads was washed, added to the samples and incubated for 1 h at 4°C. Beads were again removed by magnet. Proteins were eluted by incubating in 30μl of 0.5mg/ml Influenza Hemagglutinin (HA) Peptide (Stratech Scientific) dissolved in elution buffer (0.1M Tris pH 7.4, 150mM NaCl, 0.1% SDS, 0.5% NP40) and 1μl of 0.1M DTT (Invitrogen) was added to samples. Eluted protein samples were boiled in 4x sample loading buffer (Invitrogen) for 10 mins at 90°C. Samples were loaded onto a 4-12% polyacrylamide gel (BioRad) and electrophoresed at 150V for 10 mins, until the sample had run through the stacking wells.

Protein-containing gel was excised and cut into 1mm^2^ pieces, destained, reduced using DTT, alkylated using iodoacetamide and subjected to enzymatic digestion with sequencing grade trypsin (Promega, Madison, WI, USA) overnight at 37°C. After digestion, the supernatant was pipetted into a sample vial and loaded onto an autosampler for automated LC-MS/MS analysis.

LC-MS/MS experiments were performed using a Dionex Ultimate 3000 RSLC nanoUPLC system (Thermo Fisher Scientific) and a Q Exactive Orbitrap mass spectrometer (Thermo Fisher Scientific). Separation of peptides was performed by reverse-phase chromatography at a flow rate of 300 nL/min and a Thermo Scientific reverse-phase nano Easy-spray column (Thermo Scientific PepMap C18, 2μm particle size, 100A pore size, 75 μm i.d. × 50 cm length). Peptides were loaded onto a pre-column (Thermo Scientific PepMap 100 C18, 5μm particle size, 100A pore size, 300μm i.d. × 5mm length) from the Ultimate 3000 autosampler with 0.1% formic acid for 3 mins at a flow rate of 15μL/min. After this period, the column valve was switched to allow elution of peptides from the pre-column onto the analytical column. Solvent A was water + 0.1% formic acid and solvent B was 80% acetonitrile, 20% water + 0.1% formic acid. The linear gradient employed was 2-40% B in 90 mins (the total run time including column washing and re-equilibration was 120 mins).

The LC eluant was sprayed into the mass spectrometer by means of an Easy-spray source (Thermo Fisher Scientific Inc.). All *m/z* values of eluting ions were measured in an Orbitrap mass analyzer, set at a resolution of 35000 and scanned between *m/z* 380-1500. Data dependent scans (Top 20) were employed to automatically isolate and generate fragment ions by higher energy collisional dissociation (HCD, Normalised collision energy (NCE):25%) in the HCD collision cell and measurement of the resulting fragment ions was performed in the Orbitrap analyser, set at a resolution of 17500. Singly charged ions and ions with unassigned charge states were excluded from being selected for MS/MS and a dynamic exclusion of 60 seconds was employed.

Post-run, all MS/MS data were converted to mgf files and the files were then submitted to the Mascot search algorithm (Matrix Science, London UK, version 2.6.0) and searched against a common contaminants database (125 sequences; 41129 residues); and the CCP_*Plasmodium_falciparum Plasmodium_falciparum*_20190315 (5449 sequences; 4173922 residues) database. Variable modifications of oxidation (M) and deamidation (NQ) were applied as well a fixed modification of carbamidomethyl (C). The peptide and fragment mass tolerances were set to 20ppm and 0.1 Da, respectively. A significance threshold value of p<0.05 and a peptide cut-off score of 20 were also applied.

Data were then analysed using MaxQuant software version 1.6.17.0 (Elias and Gygi, 2007). Files were searched against *Plasmodium falciparum* 3D7 PlasmoDB-50 annotated proteins database (downloaded February 2021). Protein N-terminal acetyl and methionine oxidation were set as variable modifications, whilst cysteine carbidomethylation was a fixed modifier. C-terminal arginine was set as the enzyme specificity and trypsin as the protease. Minimum peptide length was 7 amino acids and maximum for peptide recognition was 4600 Da.

### GO enrichment analysis

The analysis tool in PlasmoDB (Aurrecoechea et al., 2009) was used to obtain GO terms for all gene IDs encoding proteins found by co-immunoprecipitation. Enrichment of GO terms versus their representation in the whole genome was calculated within PlasmoDB, with a cutoff of p=0.05 for statistically significant enrichment. Correction for multiple comparisons was conducted by both Benjamini-Hochberg FDR and the more stringent Bonferroni method, and GO terms with p-values remaining below 0.05 were considered to be enriched.

## Supplementary Figures and Tables

**S1 Figure.**
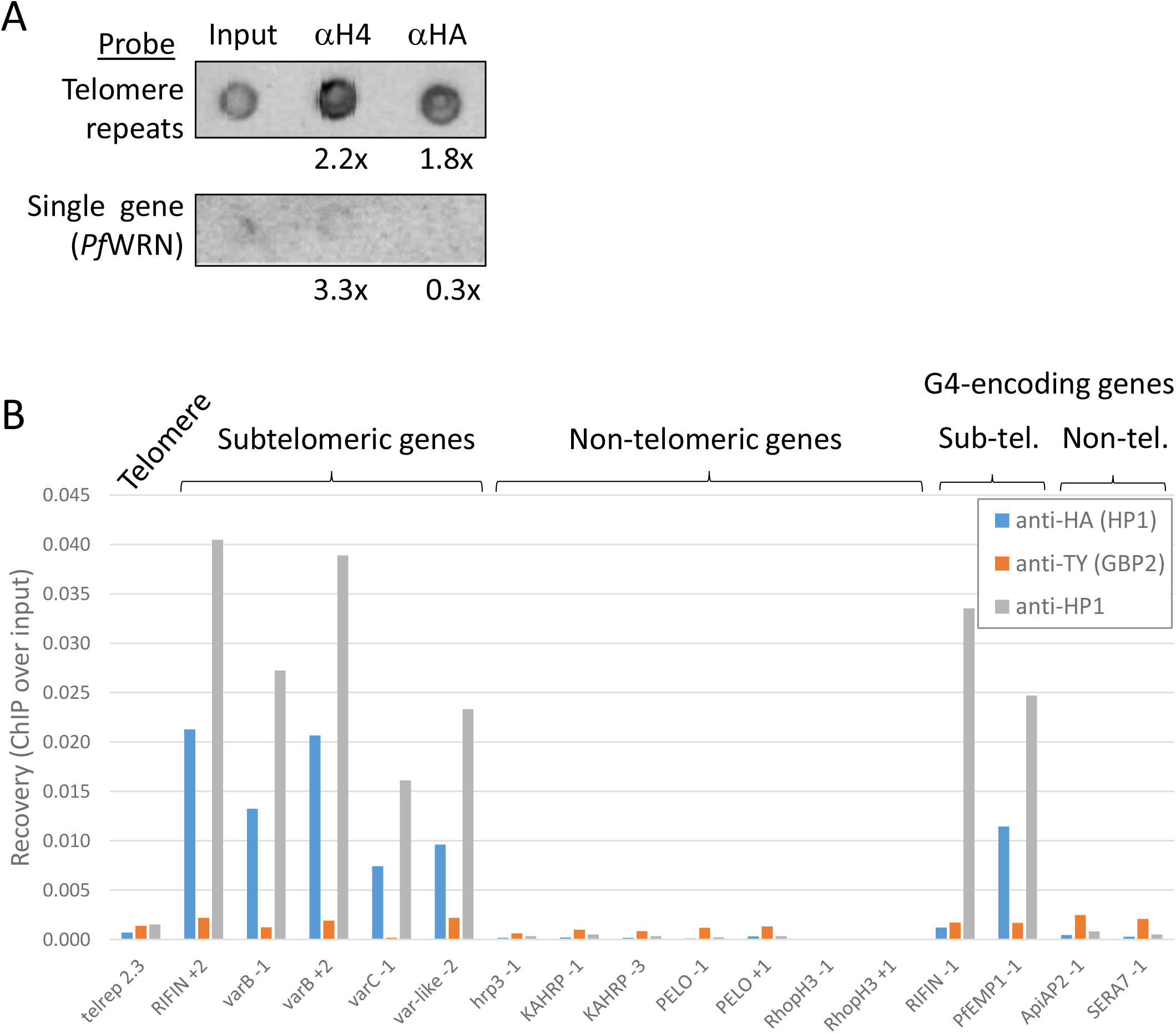
ChIP on *Pf*GBP2. (A) Dotblots of chromatin immunoprecipitated with antibodies to *Pf*GBP2-3HA or to histone H4. Fold enrichment over input is shown. Telomere sequence is enriched by ∼2x with antibody to *Pf*GBP2-HA as well as antibody to histone H4, whereas a control non-telomeric gene is detected only in histone ChIP and is not enriched in HA ChIP. (B) Examples of ChIP data from the *Pf*GBP2-Ty line which co-expresses *Pf*HP1-HA. Genome-wide ChIP-seq conducted in duplicate on this line, and also on the *Pf*GBP2-3HA line, failed to give signals significantly above background (data not shown). To confirm the validity of the ChIP procedure using the validated chromatin protein *Pf*HP1 as a control, ChIP was conducted with a selection of individual primer pairs directed to telomere repeats, subtelomeric genes, G-quadruplex-encoding genes, or chromosome-internal genes. Representative data from one of two duplicate experiments are shown. *Pf*HP1-HA, as expected, was enriched by >50-fold at sub-telomeric loci compared to chromosome-internal loci when it was immunoprecipitated with either anti-HA-tag or anti-HP1 antibodies. By contrast, *Pf*GBP2-Ty, present in the same parasites, was not strongly enriched at any locus: the mean enrichment at sub-telomeric and G4-encoding genes compared to non-telomeric genes was only 2.4-fold.

**S1 Table. Protein sequence matching data from FASP and GASP analysis of PICh extracts**.

Control (Ctl) and Target data from FASP (experiment 1) are compared to a second dataset obtained via GASP (experiment 2) using Scaffold, with a 95% threshold for proteins and peptides, and a minimum of two peptides/hit. ‘Exclusive Unique Peptide Count’ is shown for each sample. Searches were performed using Mascot against a hybrid database containing sequences derived from *Plasmodium falciparum* (download date 20^th^ July 2015), combined with common contaminant proteins (cRAP, download date 30^th^ January 2015 and contaminants, download date 13^th^ July 2012, both downloaded from Matrix Science). No *Plasmodium* peptides were observed in the control GASP experiment, possibly due to a technical error, although they were detected in the preceding FASP experiment.

**S2 Table. Table of oligonucleotide sequences**

In PICh probes, ‘C18’ refers to the flexible linker region. Locked Nucleic Acid (LNA) bases are in capitals. ‘y’ is a 67% T, 33% C custom base mix, designed to mimic the variable position in the *Plasmodium* telomere repeat (GGGTT(T/C)A) where the ratio is approx. 67% T: 33%C.

**S3 Table. Proteins immunoprecipitated with *Pf*GBP2**

Tab 1 shows all protein hits from duplicate anti-HA immunoprecipitations of *Pf*GBP2-3HA, after any proteins also found in a control IP experiment from wildtype parasites have been screened out. Tab 2 shows the subset of proteins that were reproducibly found in both *Pf*GBP2-3HA immunoprecipitations. Tab 3 shows proteins found uniquely in the control IP. GO terms for each hit are listed in the classes ‘cellular component’, ‘biological function’ and ‘molecular process’.

**S4 Table. GO terms enriched amongst proteins immunoprecipitated with *Pf*GBP2**

Enriched GO terms in the classes ‘cellular component’, ‘biological function’ and ‘molecular process’ are shown for the reproducible *Pf*GBP2 interactors, and also for total set of *Pf*GBP2 interactors found in at least one IP experiment. GO terms with statistically significant enrichment after correction for multiple comparisons are highlighted in yellow. Terms associated with RNA are in red text, and with DNA in blue text.

## Notes

### Competing Interest Statement

The authors have declared no competing interest.

